# Polinton-like Viruses Associated with Entomopoxviruses Provide Insight into Replicon Evolution

**DOI:** 10.1101/2023.10.16.562556

**Authors:** Zachary K. Barth, Ian Hicklin, Julien Thézé, Jun Takatsuka, Madoka Nakai, Elisabeth A. Herniou, Anne M. Brown, Frank O. Aylward

## Abstract

Polinton-like viruses (PLVs) are a diverse group of small integrative dsDNA viruses that infect diverse eukaryotic hosts. Many PLVs are hypothesized to parasitize viruses in the phylum *Nucleocytoviricota* for their own propagation and spread. Here we analyze the genomes of novel PLVs associated with the occlusion bodies of entomopoxvirus (EV) infections of two separate lepidopteran hosts. The presence of these elements within EV occlusion bodies suggests that they are the first known hyperparasites of poxviruses. We find that these PLVs belong to two distinct lineages that are highly diverged from known PLVs. These PLVs possess mosaic genomes, and some essential genes share homology with mobile genes within EVs. Based on this homology and observed PLV mosaicism, we propose a mechanism to explain the turnover of PLV replication and integration genes.

## Introduction

Viral satellites are genetic hyperparasites, i.e. parasites of other parasites, that rely on host or ‘helper’ viruses for their own propagation and spread (Gnanasekaran and Chakraborty 2018). Viral satellites are found throughout the tree of life, associating with viruses that infect each domain (de Sousa et al. 2023; Ren et al. 2013; Barreat and Katzourakis 2023). Within eukaryotes, satellites have been described in diverse hosts, including humans (Taylor 2012; Meier et al. 2020), bees (Olivier et al. 2008), yeast (Vepštaitė-Monstavičė et al. 2018), amoebae (La Scola et al. 2008), algae (Santini et al. 2013), and many plant species (Fritsch and Mayo 2018).

Historically, recognized eukaryotic satellites have been small entities, with RNA or single stranded DNA genomes less than 6kb, and few to no coding sequences (Gnanasekaran and Chakraborty 2018). This trend has been challenged in recent years with the discovery of hyperparasitic dsDNA viruses that integrate into eukaryotic cell chromosomes, and have genomes in the range of ∼15-25kb. These viruses have been nicknamed ‘virophages’ because of their lifestyle where they ‘infect’ and parasitize the viral factories of nucleocytoviruses (phylum *Nucleocytoviricota*) (Fischer 2021). To date, culture systems have been established for six of these virophages (La Scola et al. 2008; Fischer and Suttle 2011; Gaia et al. 2014; Hauröder et al. 2018; Sheng et al. 2022; Roitman et al. 2023). Five of these established systems belong to a monophyletic clade classified as the family *Lavidaviridae* (Fischer 2021). A culture system has also been established for a satellite that has virophage-like life cycle, but is not part of *Lavidaviridae*. This element, PgVV Gezel 14-T, parasitizes Phaeocystis globose virus (Roitman et al. 2023), and belongs to a large family of polinton-like viruses (Bellas et al. 2023).

<u>P</u>olinton-like <u>v</u>iruses (PLVs) are capsid encoding elements related to the maverick-polinton class of self-synthesizing transposons (Koonin and Krupovic 2017). Aside from PgVV Gezel 14-T, a single PLV, TsV-N1, has been experimentally characterized. TsV-N1 was found to cause autonomous lytic infection in the alga *Tetraselmis striata*(Pagarete et al. 2015). Despite the paucity of experimental PLV systems, PLV genomes are abundant in metagenomic samples and as integrated proviruses within cellular genomes (Bellas and Sommaruga 2021; Barreat and Katzourakis 2021; Bellas et al. 2023); PLVs are found in diverse eukaryotic genomes, but are notably absent in those of mammals and land plants(Barreat and Katzourakis 2021). There has also been a report of mobilization and replication of two PLVs in lepidopteran cell lines used to produce recombinant baculovirus virus-like particles (Starrett et al. 2021). While some PLVs, including TsV-N1, have genomes in the 30-40kb range, overlapping with the autonomous adenoviruses, many are closer in size to virophages. In addition to the small size of many PLVs, the virophage Mavirus has several genes with homologs found in PLVs, leading some to suggest PLVs evolved from a virophage ancestor (Fischer and Suttle 2011), and that PLVs may retain antagonism towards nucleocytoviruses as a conserved trait (Barreat and Katzourakis 2021).

Here we have analyzed the genomes of three novel PLVs isolated from entomopoxvirus derived occlusion bodies formed during infection of two lepidopteran host species (Thézé et al. 2013). The association of these PLVs with entomopoxvirus occlusion bodies provides compelling evidence for these elements possessing a hyperparasitic lifestyle and is the first indication of a virophage-like element associated with poxviruses. Aligned with this hypothesis, we observe evidence of horizontal gene transfer between these PLVs and entomopoxviruses. Among these entomopoxvirus associated PLVs, we see two distinct lineages based on phylogenies of structural gene modules. These lineages encode divergent replication modules and possess capsid genes highly diverged from other PLV lineages, demonstrating previously unrecognized diversity amongst eukaryotic small dsDNA viruses. We also observe replicon gene turnover amongst related elements and propose a mechanism to explain these patterns.

## Results and Discussion

### Lepidopteran entomopoxvirus associated elements resemble PLVs

We set out to analyze the genomes of three elements that had been previously isolated from the occlusion bodies of lepidopteran infecting entomopoxviruses (EVs)(Thézé et al. 2013). Two elements were associated with Adoxophyes honmai entomopoxvirus (AHEV), and one element was associated with Choristoneura biennis entomopoxvirus (CBEV). The genomes had been submitted into NCBI with the designation ‘entomopoxvirus virophages’ as their occurrence within EV occlusion bodies was highly suggestive that these elements parasitized EVs. While annotating these genomes, we found that they possessed several signature genes for polinton-like viruses (PLVs) and opted to rename them accordingly to AHEV_PLV1, AHEV_PLV2, and CBEV_PLV1. Additionally, through a tBLASTn search, we were able to identify an element into chromosome 9 of the coleopteran *Cetonia aurata* that bore substantial homology to some proteins encoded by AHEV_PLV2. We named this fourth element Ca_PLV1. All of the elements contained detectable terminal repeats, indicating that they are complete.

Using HHPRED to predict gene functions, we found that the four novel PLVs encoded replication machinery characteristic of PLVs: AHEV_PLV1 and Ca_PLV1 each encode a family B DNA polymerase (PolB), while AHEV_PLV2 and CBEV_PLV1 encode a gene with a C-terminal D5-like SF3 helicase domain (Figure 1). While HHPRED was not able to detect a primase domain for these genes, we hypothesize that they are primase-helicases with a primase-polymerase domain, as this arrangement is common in PLV genomes(Yutin et al. 2015), and a primase-polymerase domain would presumably be necessary to support replication. As with many PLVs(Yutin et al. 2013; Yutin et al. 2015; Koonin and Krupovic 2017), the putative AHEV_PLV2 and CBEV_PLV1 primase-helicases are accompanied by a tyrosine-recombinase.

**Figure 1.**
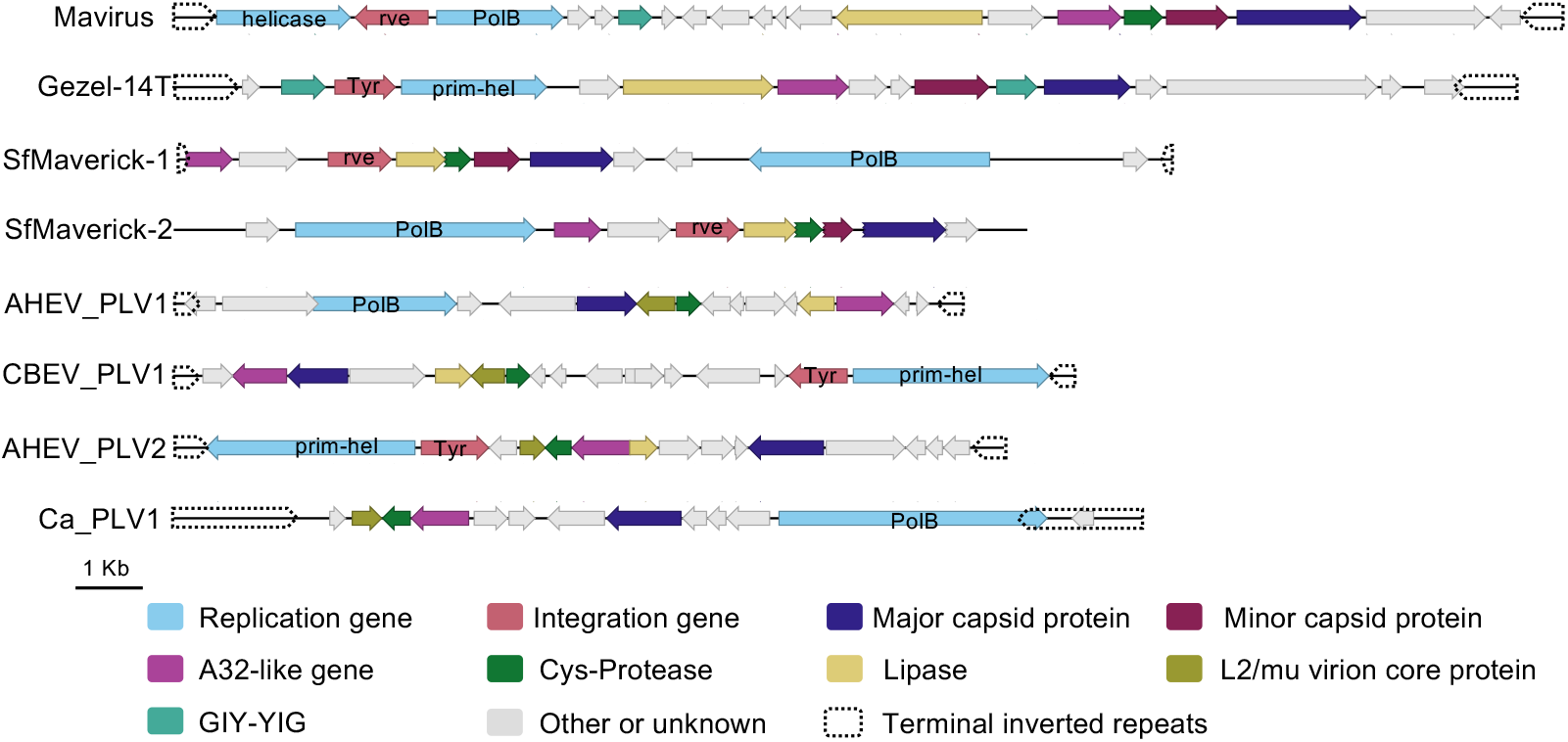
Entomopoxvirus associated elements possess gene content characteristic of PLVs. Diagrams of several PLVs and the PLV-like virophage Mavirus. Functional gene classes and TIRs are denoted by color. Replication and integration genes are labeled. ‘Tyr’ is tyrosine recombinase. ‘Prim-hel’ is primase-helicase. The genomes of AHEV_PLV1, CBEV_PLV1, and AHEV_PLV2 were assembled as circular genomes and have been linearized at their terminal inverted repeats for ease of comparison.

Aside from PolB, one of the signature genes of the maverick-polinton group of PLVs is a retroviral-type integrase (rve) gene. An example of this can be seen with SfMaverick1 and 2, from the lepidopteran species *Spodoptera frugiperda* (Figure 1). Curiously, we were unable to detect any integration genes within AHEV_PLV1 and Ca_PLV1. As AHEV_PLV1 was found in occlusion bodies, it may be non-integrative; however, Ca_PLV1 was found integrated within the *C. aurata* genome. It is possible that these elements possess cryptic integration functions unknown to current gene annotations, or they may rely on integration genes encoded *in trans*. Regardless, the lack of a detectable integrase does not appear to preclude the possibility of integration by these elements.

In addition to encoding replication genes characteristic of PLVs, the four novel elements encoded multiple morphogenesis genes associated with PLVs and virophages. All four elements encoded an A32-like genome packaging protein, and an Adenovirus-type cysteine-protease, and the three lepidopteran elements encoded a protein similar to the phospholipase domain of parvovirus virion protein VP1 (Figure 1). These three gene classes have been described in PLVs and virophages(Yutin et al. 2013; Yutin et al. 2015; Koonin and Krupovic 2017). Additionally, we found that the four novel elements encoded a homolog of the adenovirus Late L2 mu core protein (Figure 1). While the precise role of this adenovirus protein is unknown, it may be involved in viral chromatin condensation for packaging(Anderson et al. 1989). To our knowledge, this protein class has not been linked to PLVs or virophages previously. It has been suggested that PLVs may comprise a sister taxa to adenoviruses due to the relatedness of their shared gene content(Barreat and Katzourakis 2023). The L2 mu core protein provides yet another example of gene classes shared between both groups.

### EV-associated PLVs have highly divergent capsid proteins

Surprisingly, HHPRED did not detect capsid proteins within the four novel elements. While we initially considered the possibility that these elements might not encode their own capsid genes, de novo structural prediction with Alphafold(Jumper et al. 2021) revealed that each element encoded proteins predicted to form jelly-roll folds characteristic of PLV capsid proteins (Figure 2, A-C)(Krupovic et al. 2014). In particular, the AHEV_PLV2 structural model was a good match for a double jelly roll-fold characteristic of major capsid proteins (MCPs) (Figure 2 B)(Krupovic et al. 2014). The candidates for AHEV_PLV1 and CBEV_PLV1 were not predicted to form a second jelly-roll fold (Figure 2 A and C), but rather consisted of a jelly roll fold link to a second beta-sheet rich domain. A slight reconfiguration of this beta-sheet region could potentially reorganize to form a jelly roll fold like structure, rendering this predicted fold and structural organization in alignment with having a structural similarity to known capsid proteins. Based on our sequence and structural predictions, these proteins are most likely the coding sequences to form major capsid proteins encoded by these elements. It is possible that the unformed jelly roll-fold, but beta strand rich region, predicted for these proteins may reflect that they are highly diverged from capsids that have resolved structures.

**Figure 2.**
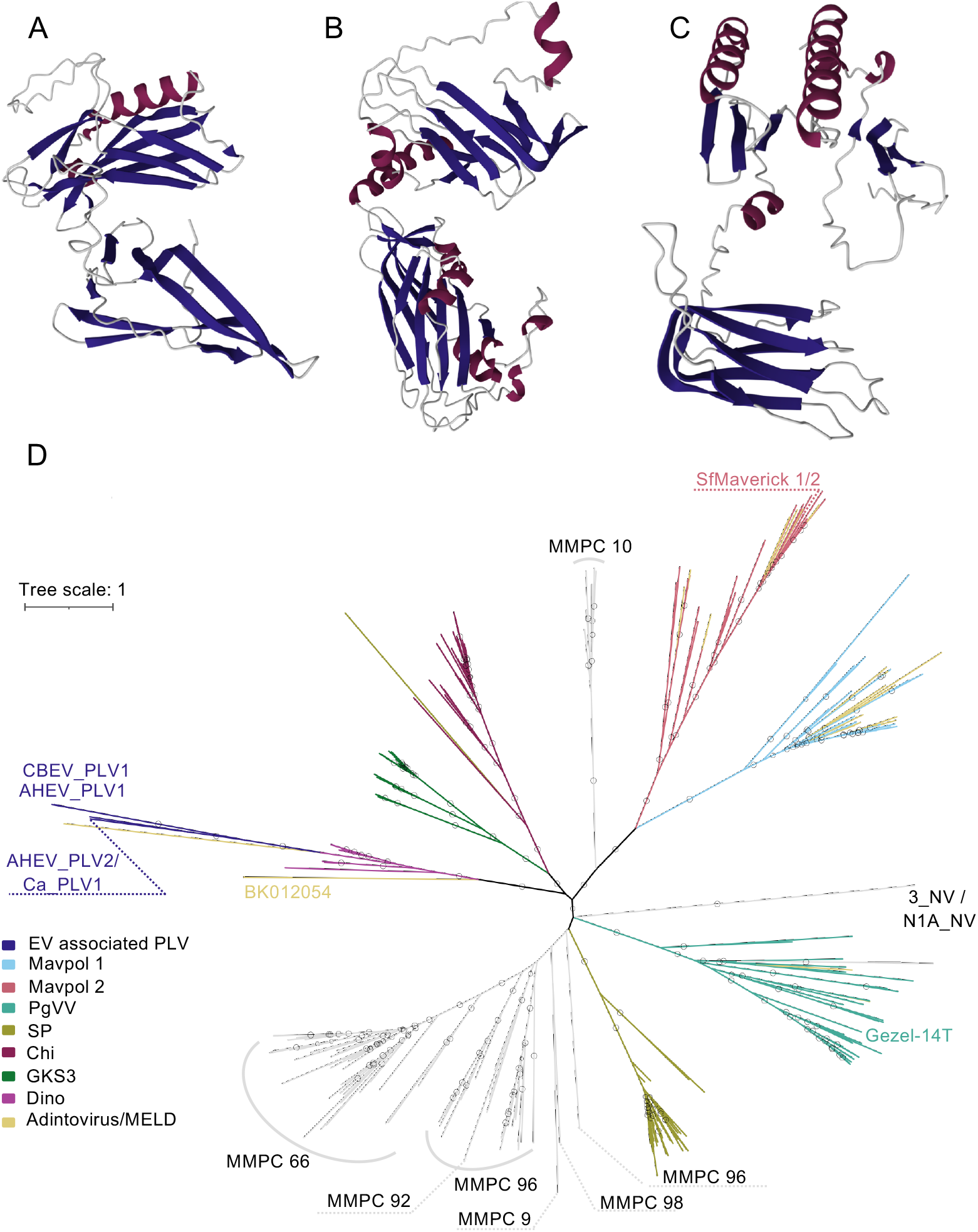
Entomopoxvirus associated PLVs have highly diverged capsid proteins. A-C, de novo structural predictions of putative major capsid proteins for AHEV_PLV1(A), AHEV_PLV2(B), and CBEV_PLV1 (C). Beta sheets are colored dark blue and helices are colored purple. D. A phylogeny of major capsid proteins genes amongst PLVs. Several major lineages of PLVs are denoted by color, and additional MCP clusters are labelled based on classification in X and Y. PLVs from figure 1 are also labelled.

Next, we built a phylogeny that included the EV associated PLV MCPs as well as MCPs from Mavpol elements and MCPs belonging to a large cluster related to PgVV MCPs that had been previously identified (Bellas et al. 2023). These clusters provide a considerable amount of breadth of cellular host species, and cover nearly all PLVs that have been found in animal species. The EV associated PLV capsid sequences clustered into a clade that also contained a MELD virus from the *Anelosimus* tangle-web spider (BK012043), suggesting the existence of a novel lineage of animal tropic PLVs. All of the MCP proteins within this clade had long branches, suggesting that they are more distantly related to known MCP groups (Figure 2D). The placement of this EV associated PLV clade was unstable when we made slight alterations to our data set. An earlier version of our data set contained three fewer MELD virus genomes, one from a *Powellomyces* chytrid fungus (BK012054) and two from algal species (BK012050 and BK012048). A tree made from this older data set placed MCPs from the cnidarian encoded 3_NV and N1A_V elements within the EV associated PLV clade, and placed this clade as a sister taxon to the SP group, which includes PLVs found in the fungus *Spizellomyces punctatus* (Bellas and Sommaruga 2021), (Figure S1). In our later tree, the EV associated PLV clade is placed within the Dinoflagellate specific clade, and is unrelated to 3_NV and N1A_V (Figure 2D). The instability of their tree topology highlights the high level of divergence that EV associated PLV MCP sequences have from those of other PLV elements. Visual inspection of the MCP multiple sequence alignment revealed that the MCPs from AHEV_PLV1 and CBEV_PLV1 were particularly poorly aligned (Data S1). Thus, only preliminary conclusions can be drawn regarding the relationship between the MCPs of EV associated PLVs and those of other PLV groups

Within the EV associated PLV clade, the AHEV_PLV1 and CBEV_PLV1 MCPs were closely related to each other (Figure 2D). The AHEV_PLV2 and Ca_PLV1 MCPs were also closely related to each other, but highly divergent from those of AHEV_PLV1 and CBEV_PLV1 (Figure 2D). In summary, our tree suggests that the EV associated PLV MCPs exist as two distinct lineages within a highly diverged group of animal PLVs. This grouping, along with the highly derived nature of EV associated PLV capsids, shows the potential wealth of PLV diversity that remains undiscovered.

### EV associated PLVs are mosaic elements

The close relationships of MCPs between AHEV_PLV1 and CBEV_PLV1, and AHEV_PLV2 and Ca_PLV1, respectively, was intriguing given that AHEV_PLV1 and Ca_PLV1 encode PolB replicases, while AHEV_PLV2 and CBEV_PLV1 encode a primase-helicase. Using BLASTp, we looked for the presence of other homologous genes amongst the novel PLVs. The independent clustering of capsids for AHEV_PLV1 with CBEV_PLV1, and AHEV_PLV2 with Ca_PLV1 was extended to most of the morphogenic genes we identified (Figure 3A and 3B), revealing two lineages of morphogenic gene modules. Despite belonging to separate morphogenic lineages, AHEV_PLV2 and CBEV_PLV1 shared homology in their integration and replication genes (Figure 3C). Notably, homology was variable on a sub-protein level. The AHEV_PLV2 and CBEV_PLV1 primase-helicases shared 26% amino acid identity across a 312aa n-terminal region (Figure 3C), while the rest of the proteins, including the helicase domains, did not show detectable similarity, suggesting the prior occurrence of sub-genic domain swapping. Taken together, these observations show that EV associated PLVs are mosaic elements and have exchanged gene modules and functional domains over the course of their evolution.

**Figure 3.**
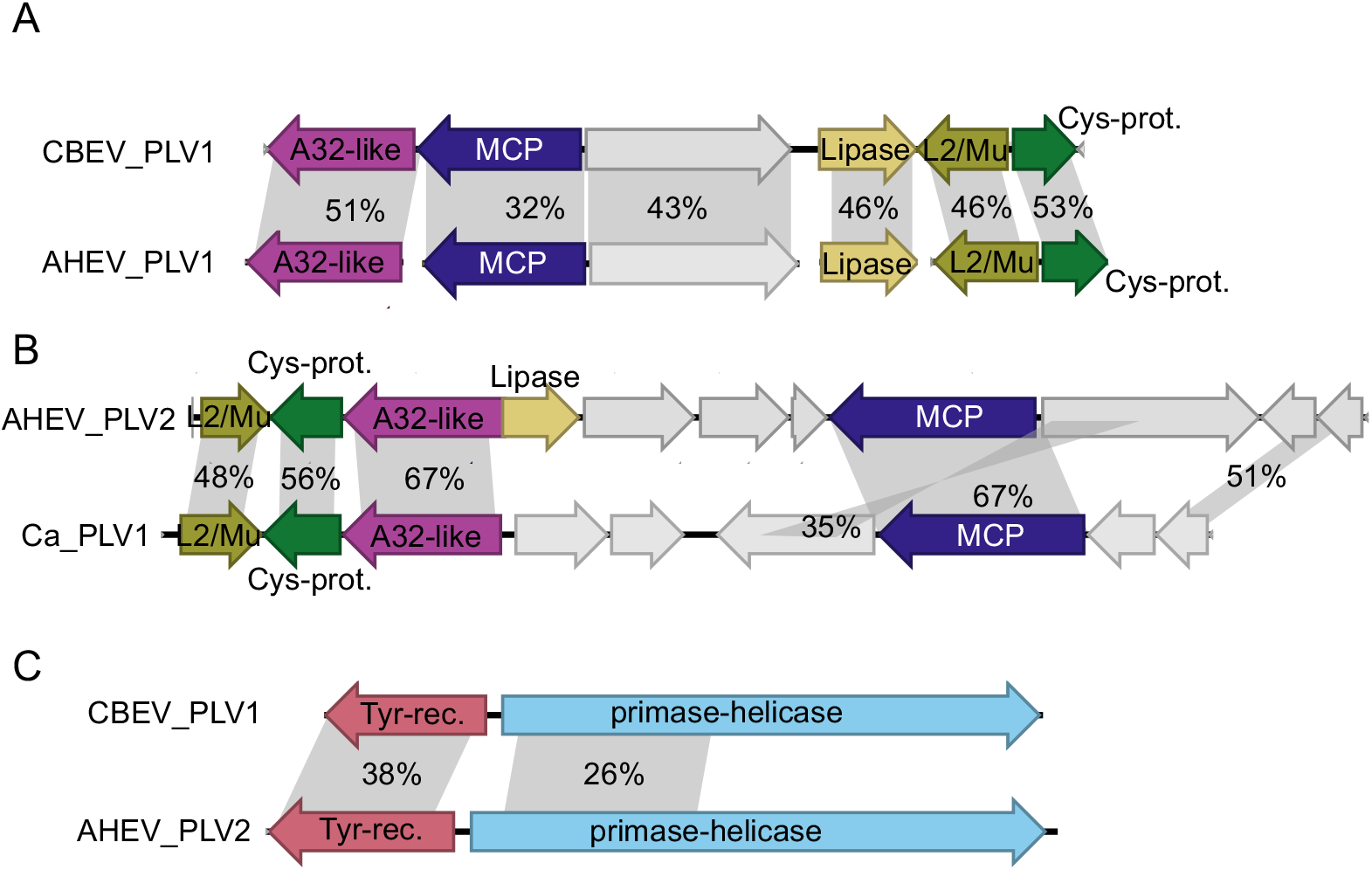
Entomopoxvirus associated PLVs exhibit genomic mosaicism. Comparisons of entomopoxvirus associated PLV loci with percent amino acid identity indicated across the shaded regions for CBEV_PLV1 and AHEV_PLV1 structural genes (A), AHEV_PLV2 and Ca_PLV1 structural genes (B), and CBEV_PLV1 and AHEV_PLV2 integration and replication genes (C).

### An EV-associated PLV end module is related to EV-encoded genes

It may be possible to infer aspects of PLV biology from their genomic organization. For both AHEV_PLV2 and CBEV_PLV1, the primase-helicase gene is encoded between one of the terminal inverted repeats (TIRs) that flank the element, and the tyrosine recombinase. Tyrosine recombinases typically use inverted repeats as their substrates for recombination (Rajeev et al. 2009), and it is common within integrating elements for the integrase to be encoded next to the integration site (Smyshlyaev et al. 2021). The placement of the primase-helicase between the TIRs and tyrosine recombinase suggests these three PLV components may interact, with the TIRs functioning both as a substrate for integration by the tyrosine recombinase, and as the origin of replication recognized by the primase-helicase. We hypothesize that such cis-acting functions in the TIRs would select for co-clustering of their interacting proteins, giving rise to functional modules at the end of the genome. Consistent with this replicative end-module hypothesis, many previously published PLV genomes display replication and integration genes encoded near each other at one end of the element, or the replication and integration genes encoded at opposite ends of the element(Yutin et al. 2013; Yutin et al. 2015; Koonin and Krupovic 2017). Indeed, the replicases of all four of the novel PLVs described here are located near one of the TIRs (Figure 1).

Further supporting the existence of a replicative end module, we found that all three components of CBEV_PLV1’s end module had homologs within the genome of Mythimna separata entomopoxvirus (MySEV) (Figure 4). Within the CBEV_PLV1 and MySEV TIRs, an approximately 300bp region with 72% nucleotide identity occurs less than 300bp from the genomic termini (Figure 4B). Similarly, MySEV encoded a tyrosine recombinase (Figure 4C) and a primase-helicase (Figure 4D) that were closely related to those of CBEV_PLV1. While these MySEV components were encoded in separate regions of the MySEV genome, poxviruses have been hypothesized to initiate replication out of hairpin structures, referred to as telomeres, that occur within the TIRs(Shenouda et al. 2022). Tyrosine recombinase activity on these ends could be a mechanism for genome dimer resolution, as occurs in bacterial genomes, plasmids, and some bacteriophages (Castillo et al. 2017; Haenebalcke and Haigh 2013).

**Figure 4.**
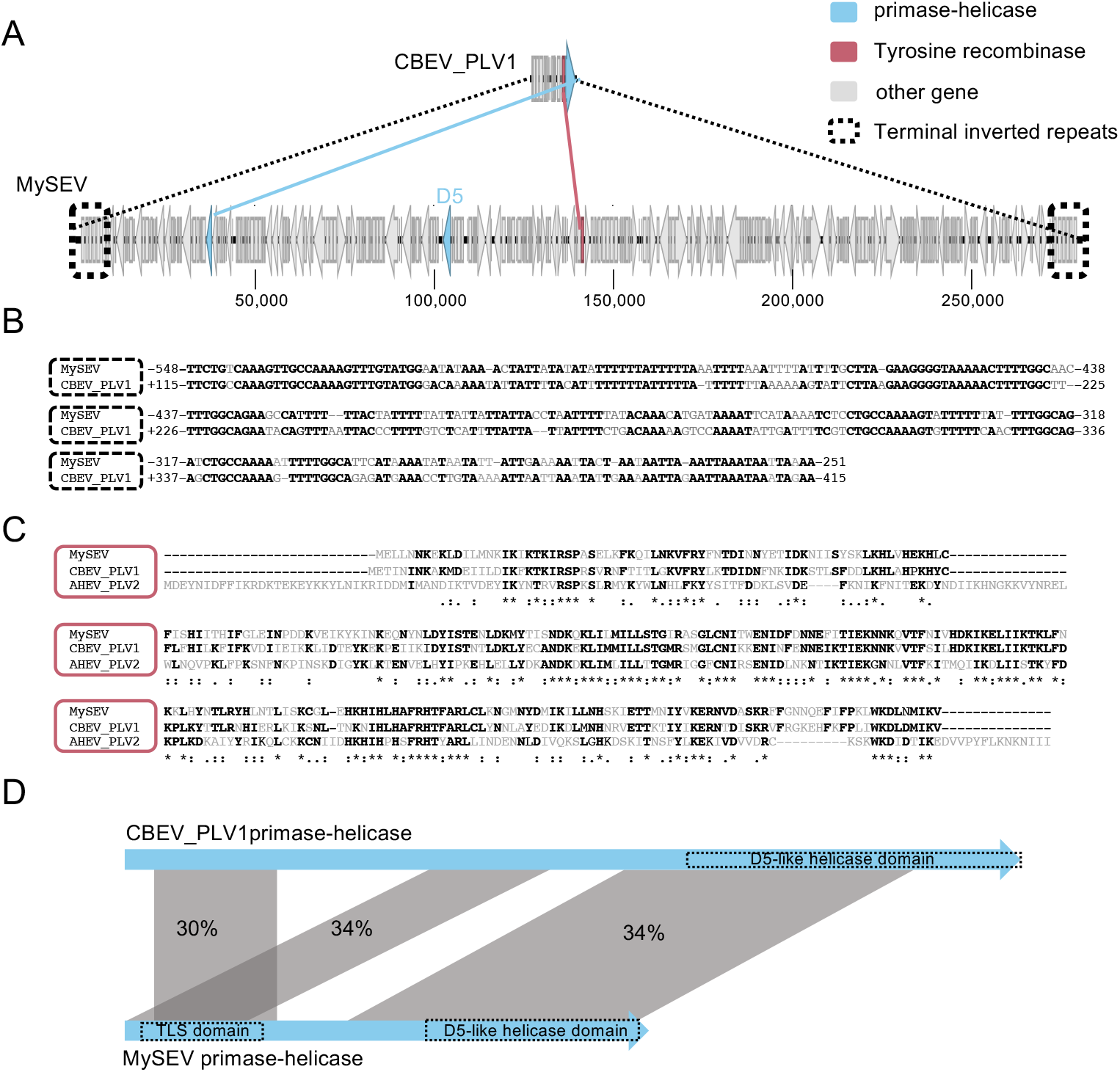
PLV end module shows homology to entomopoxvirus genes and non-coding regions. A. Diagram of CBEV_PLV1 and MySEV genomes with homologous features labeled and indicated by colored lines. B. An alignment of homologous nucleotide subsequences that occur within the TIRs of CBEV_PLV1 and MySEV. Mismatched bases are grey, matching bases are black and bolded. C. An alignment of the tyrosine recombinase protein sequence from MySEV, CBEV_PLV1 and AHEV_PLV2. Positions with a consensus across at least two of the three proteins are bolded in black. ‘.’ indicates conservation across groups with weakly similar properties. ‘:’ indicates conservation across groups with strongly similar properties. ‘*’ indicates complete conservation of the residue. D. Diagram showing the shared amino acid identity across shaded regions for MySEV and CBEV_PLV1 primase-helicase genes.

Notably, HHPRED predicted similarity to the human primase-polymerase primpol in an N-terminal region of the MySEV primase-helicase (Figure 4D) that bore shared sequence identity with the CBEV_PLV1 primase-helicase. This region of the CBEV_PLV1 primase-helicase is also homologous to that of AHEV_PLV2 (Figure 3C), further supporting the hypothesis that the AHEV_PLV2 and CBEV_PLV1 genes do possess primase-polymerase activity and the potential for replicating the PLV genome.

The dispersed loci of poxvirus replication genes contrast with the modular architecture of the PLVs. This difference may reflect different organizational principles of PLV and poxvirus genomes. Our analysis suggests a high level of mosaicism amongst EV associated PLVs. This requires that cooperating components of functional modules be encoded together, to reduce the chance that they be separated through recombination and rendered nonfunctional. While poxviruses are prone to recombination (Bobay and Ochman 2018; Sasani et al. 2018), they have much larger and more complex genomes, and do not exhibit the same mosaic modularity as the EV associated PLVs. Modularity may be a trait of small genomes, as greater complexity would likely require greater functional integration of a genome’s components.

Like other poxviruses, MySEV encodes a D5 primase-helicase homolog which would be expected to serve as the main primase for genome replication (Figure 4A). It is unclear why MySEV would encode two primase-helicase genes. It is possible that the two primase-helicases have specialized functions. We hypothesize that the divergent primase-helicase primes replication in the TIRs. A possible function for the canonical D5 homolog then would be priming of DNA synthesis on the lagging strand.

The similarity of these MySEV components to the CBEV_PLV1 end module is remarkable in that they represent the strongest instances of both nucleotide and amino acid sequence similarity that we were able to detect between the EV associated PLVs and EVs. This high level of homology indicates relatively recent horizontal gene transfer between EVs and EV associated PLVs, and suggests that essential PLV replication and integration genes may have originated in viruses parasitized by PLVs.

### EV-associated PLVs encode replication genes related to mobile EV helicases

In addition to the MySEV encoded primase-helicase, a number of nucleocytovirus-encoded helicases shared sequence identity with the primase-helicases of CBEV_PLV1 and AHEV_PLV2 (Figure 5A and 5B). In particular, the primase-helicase from CBEV_PLV1 had sequence similarity to helicases from its co-isolated virus CBEV and CBEV’s close relative CREV, consistent with HGT between EVs and associated PLVs. In contrast, the helicase domain of AHEV_PLV2 resembled those of primase-helicases from ascoviruses, namely the lepidopteran infecting Heliothis virescens ascoviruses (HVAV) and Diadromus pulchellus ascovirus (DPAV), isolated from a parasitoid wasp that preys on lepidopterans. The relationship between the AHEV_PLV2 helicase domain and those of the two ascoviruses suggests that EV associated PLVs are part of a broader gene flow network that includes non-poxvirus nucleocytovirus genomes, and may even extend to the viruses of their own hosts’ parasitoids.

**Figure 5.**
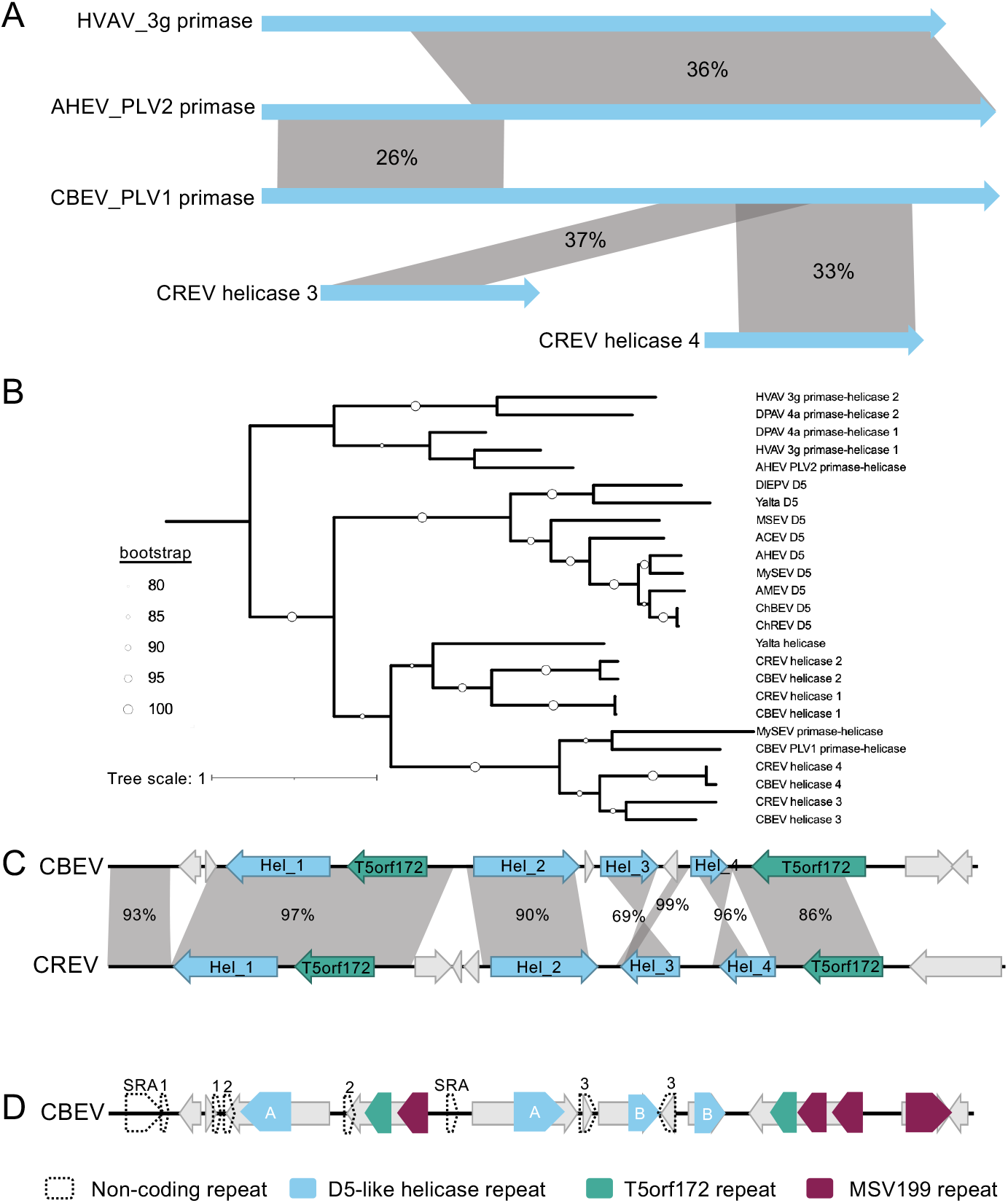
PLV replication genes are related to mobile primase-helicase genes from viruses. A. Diagram showing regions of homology across primase-helicase and helicase genes. Genes are depicted as blue arrows. Percent amino acid identity across shaded regions is indicated. B. Pairwise Identity matrix or tree of the D5-like helicase domains from proteins encoded by EVs, EV associated PLVs, and ascoviruses. C. Diagram of CBEV and CREV tandem inverted repeat (TIR) subregions. Gray shaded regions indicate areas of nucleotide similarity. ‘Hel’ is helicase. D-Repeat sequences within the CBEV TIR. Repeat types are denoted by color. For helicase-coding and non-coding repeats, homologous sequences are given the same label. ‘SRA’ is short repeat array.

All of the helicase domains with detectable similarity to the AHEV_PLV2 and CBEV_PLV1 domains have detectable similarity to the vaccinia virus primase-helicase protein D5 via HHPRED, or have close homologs where such similarity was readily detectable. Notably, these domains clustered independently from those of the core D5 homologs present within sequenced EVs (Figure 5B). In the case of the ascovirus specific cluster, the HVAV and DPAV lack a canonical D5 homolog and each have two copies of these divergent D5-like primase helicase.

D5 is considered a core gene of nucleocytoviruses, so it is likely that one or both of the primase-helicases within the ascovirus genomes are essential, but further empirical evidence is required to to infer their functions. On the other hand, all EVs possess identifiable core D5 homologs, suggesting that their divergent helicases have a distinct function.

All of the CBEV, CREV, and Yalta virus encoded helicases occur in the TIRs. Additionally, we hypothesized that the MySEV primase-helicase may act on MySEV’s TIRs due to similarities with the CBEV_PLV1 end module, further tying this helicase cluster to EV TIRs. Within nucleocytoviruses, the TIRs are often hotspots for diversity and repetitive sequences(Mönttinen et al. 2021). For example, CBEV and CREV have highly similar genomes with the bulk of their unrelated sequence being located within the TIRs (Thézé et al. 2013). A closer examination of the CBEV, CREV, and Yalta virus helicases suggested that these genes may themselves be mobile. In these genomes, the accessory helicases show an association with T5orf172 genes (Figure 5C). The T5orf172 protein domain is a subfamily of the GIY-YIG endonuclease domain, and was recently proposed to comprise a widespread family of homing endonuclease genes (HEGs) in bacteriophages (Barth et al. 2023). HEGs are mobile elements that mobilize through the cleavage of distinct, yet related, rival loci that lack the HEG. The broken DNA is repaired through homologous recombination, using the HEG coding locus as a repair template, which causes the HEG to be copied into the cognate genome, facilitating the HEG’s spread (Burt and Koufopanou 2004; Stoddard 2011; Belfort and Bonocora 2014). HEGs are highly abundant in bacteriophages (Edgell et al. 2010; Barth et al. 2023), and have also been observed within nucleocytovirus genomes (Deeg et al. 2018; Mirzakhanyan and Gershon 2020).

The helicase and T5orf172 genes in CBEV and CREV have several features that suggest their mobility. The uneven level of sequence similarity and the presence of genetic rearrangements (Figure 5C) suggests recombination at these loci, which is consistent with a homing mechanism. Similar to previously recognized HEGs (Barth et al. 2023), the EV T5orf172 genes exist as multiple copies of homologous orfs with a modular and repetitive domain structure (Figure 5D). While the helicases do not display the same modularity, they too exist as repeated homologous coding sequences (Figure 5D). As another sign of mobility, the helicases are flanked by repeat sequences, a hallmark of mobile elements (Figure 5D), (Table S1).

Beyond our own observations in EVs, an association between D5-like helicases and T5orf172 genes has been previously observed throughout nucleocytoviruses (Mönttinen et al. 2021). While the association of these gene groups likely has a mechanistic basis, the nature of that basis is hard to predict. SF1B helicases of the Pif1 family provide mobility functions for the helitron and replitron classes of mobile elements(Chandler et al. 2013; Craig 2023). Helitron and replitron helicases possess an HUH single strand endonuclease domain that nicks DNA to initiate transposition through a rolling circle replication mechanism. D5-like helicases belong to the SF3 group rather than SF1B, but could potentially also facilitate transposition through a rolling circle mechanism. The rep genes of CRESS-DNA viruses consist of a HUH endonuclease fused to a SF3-helicase domain and initiates rolling circle replication of the viral genome (Zhao et al. 2019; Tarasova and Khayat 2021). The EV helicases could conceivably mobilize through the same mechanism, with the endonuclease function being conferred by a separately encoded T5orf172 gene.

The repeat sequences flanking the helicase do not encompass the T5orf172 genes, suggesting that the genes are capable of mobilizing separately, but this organization does not preclude the possibility of these genes functioning together. HEGs sometimes mobilize adjacently encoded self-splicing introns through a process called ‘collaborative homing’ (Bonocora and Shub 2009; Zeng et al. 2009). In these instances, HEGs target the intron-less allele, facilitating the co-mobilization of the intron and HEG (Zeng et al. 2009). This arrangement has been proposed to be a predecessor of the classically described intronic HEGs found in many diverse organism (Bonocora and Shub 2009). In a similar way co-mobilization of HEGs and mobile helicases could have preceded the evolution of replitron and helitron-like elements.

The presence of these helicases in EV TIRs, and their similarity to components of the CBEV_PLV1 replication end module led us to take a closer look at PLV and virophage replication machinery. Apart from their proximity to TIRs, we were able to find other similarities between some PLV and virophage primase-helicases and mobile helicases in EVs. In multiple cases, primase-helicases or PolB genes are encoded proximally to GIY-YIG endonucleases, a protein superfamily that includes T5orf172. The putative Mavirus end module consists of an rve integrase and a PolB gene. Less than 800bp from the PolB gene is a GIY-YIG endonuclease. On the other side of the Mavirus end module, a helicase occurs between the TIR and the rve integrase. This end module appears in divergent virophage genomes (Figure 6A), and there are also variant alleles of the helicase, and GIY-YIG genes, suggesting that the end module and flanking genes are mobile. Along similar lines, a GIY-YIG endonuclease occurs between the Gezel-14T end module and its TIR (Figure 6B). Reminiscent of repeat sequences flanking EV mobile helicases, a >300bp direct repeats occurs within the Gezel-14T TIR as well as immediately downstream of the end module (Figure 6B), suggesting that the GIY-YIG endonuclease, tyrosine recombinase, and primase-helicase may all be part of a mobile module. From these observations, we have developed an evolutionary model for replicative and integrative end module turnover in PLVs and virophages (Figure 7). First, an element bearing a PolB end module has one of its TIRs invaded by a mobile helicase, disrupting the TIR sequence. Acquisition of a primase domain to the helicase, and tyrosine recombinase restores replication and recombination functions to the altered element end. The ancestral PolB and rve integrase no longer confer a benefit to the element, and will eventually be lost through mutation.

**Figure 6.**
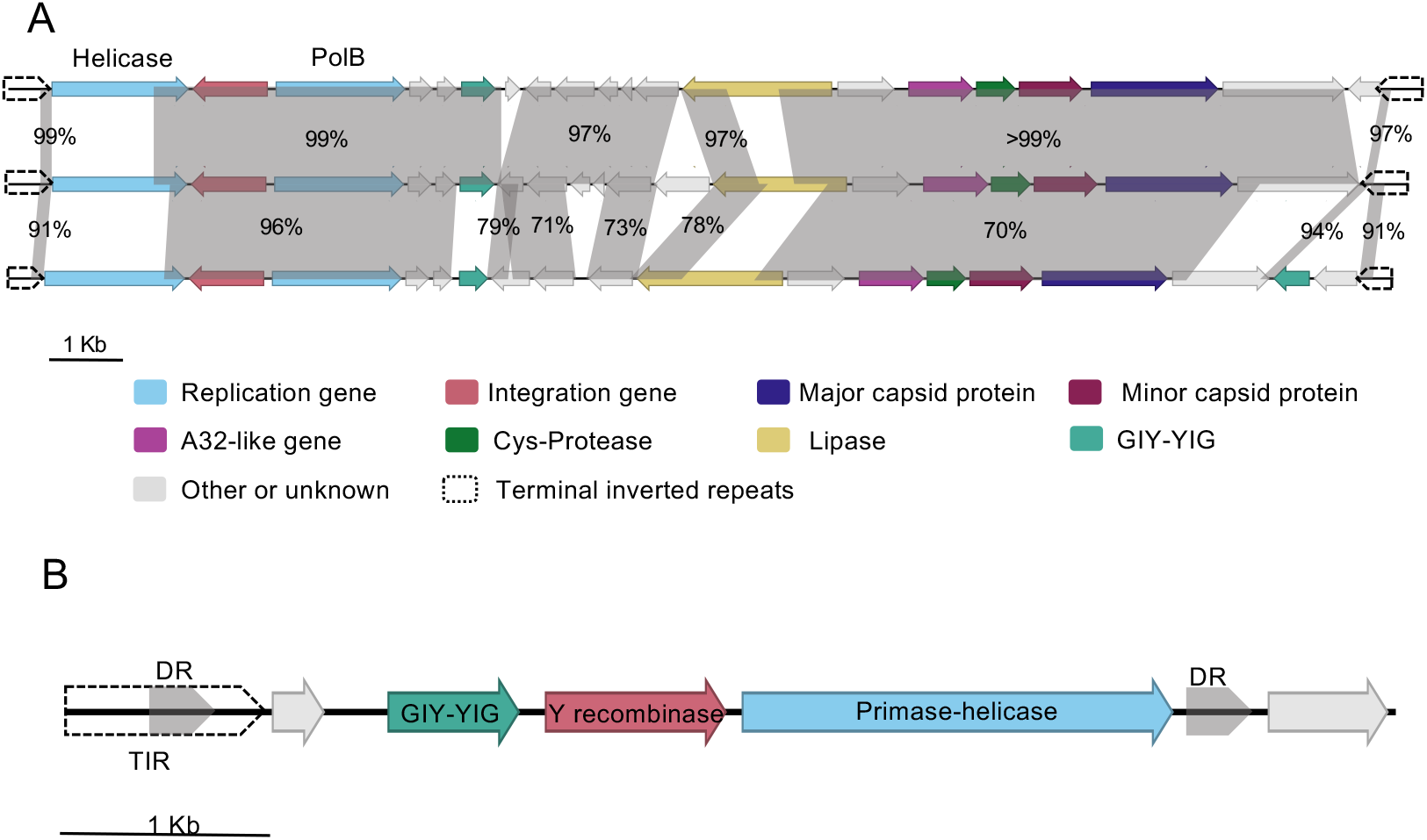
Virophage and PLV end modules bear signatures of mobility. A. Diagram showing regions of shared nucleotide identity across Mavirus isolates. TIRs and genes of interest are indicated by color. From top to bottom, Mavirus accession numbers are KU052222, OX559518, and OX560865. B. A subregion of the Gezel-14T genome with genes and genomic features labeled. ‘Y recombinase’ is tyrosine recombinase. ‘DR’ is direct repeat.

**Figure 7.**
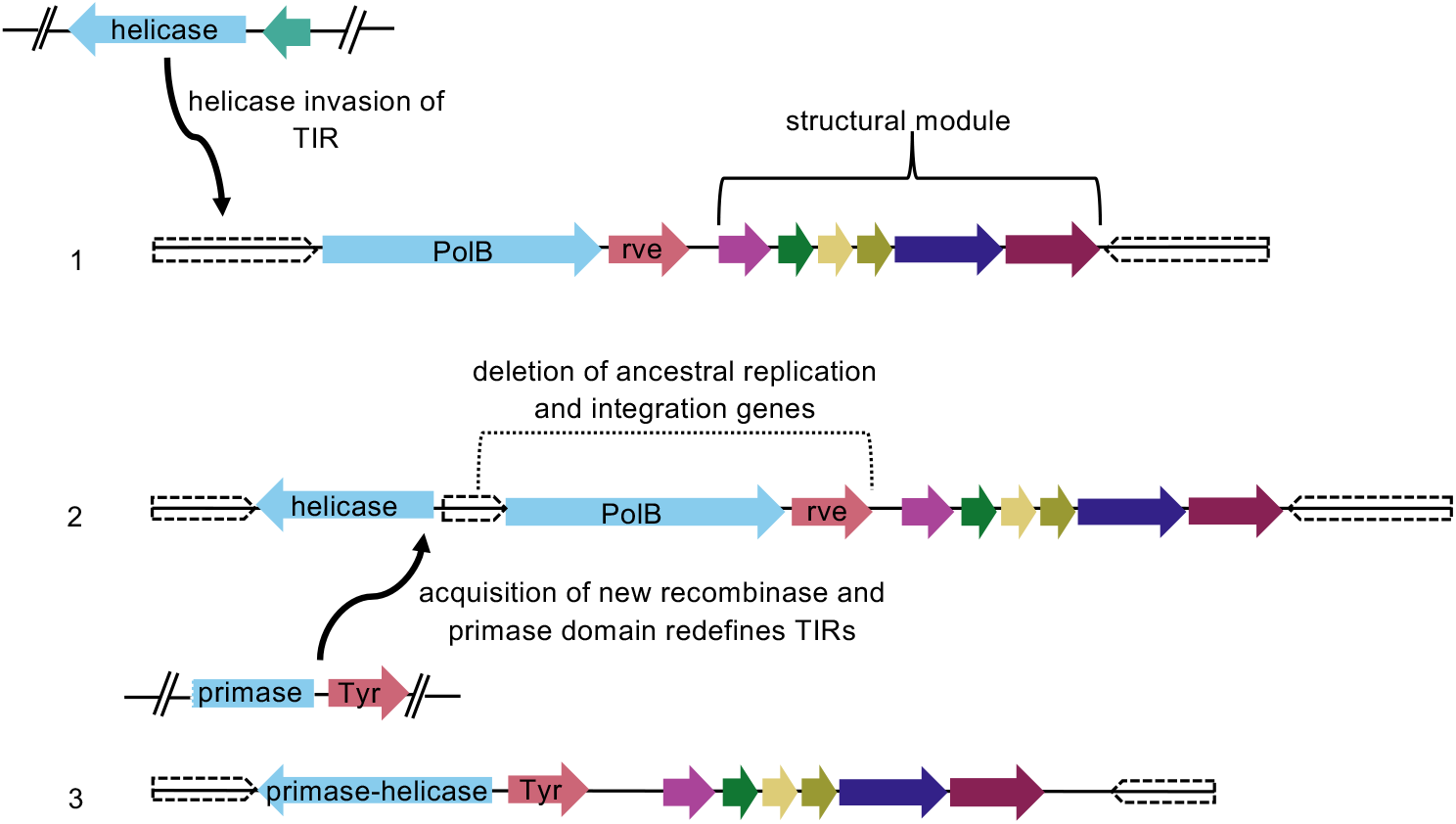
A model for PLV replicon replacement. A diagram showing a proposed mechanism for PLV replicon turnover. First, a mobile helicase gene invades the TIR of a PLV. Through recombination, the helicase gene acquires primase function and recruits a tyrosine recombinase for end maintenance, while the ancestral PLV replication and integration genes are lost through deletion.

There are many conceivable variations to this model. Recruitment of primase and recombinase functions could occur before or after invasion of the TIRs. In the case of Mavirus, we hypothesize that the Mavirus module is in an intermediate state. The rve and PolB genes are not only still present, but are presumably still providing replication and recombination functions. No additional recombinase is present, and the Mavirus helicase consists only of a helicase domain, and thus is unlikely to support replication on its own. It’s unclear if the helicase gene performs a critical replication function for Mavirus; none of the closely related virophages we found lacked a helicase, but the helicase gene was frequently substituted for alternative helicase genes. In contrast, we hypothesize that the entire Gezel-14T end module is mobile, as the endonuclease, recombinase, and replicase are all flanked by direct repeats, suggesting that they could move as a single unit. The GIY-YIG endonuclease could assist in mobility by targeting the ancestral end module. Such a homing pathway would enable end module replacement in a single step rather than relying on random mutation to delete the ancestral end module.

The validity of our end module turnover model can only become clear with experimental follow-up, but it highlights an intriguing relationship between mobile genes in nucleocytovirus TIRs, and the end modules of PLVs and virophages. Connections between chromosomal ends and mobile elements have been described in diverse systems. Some herpesviruses can integrate within the telomeres at the end of chromosomes in animals (Gennart et al. 2015; Wood et al. 2021), and more recently, some herpesvirus-related viruses were shown to integrate in the telomeres of thraustochytrid protists (Collier et al. 2023). In circular chromosomes of bacteria, many mobile elements integrate into the chromosome dimer resolution site located near the replication terminus (Midonet and Barre 2014; Midonet and Barre 2016). In line with our model, there have been multiple instances of mobile element domestication for chromosome end maintenance. In *Drosophila* telomeres are maintained by retrotransposons that integrate specifically at the ends of chromosomes (Mason et al. 2008; Pardue and DeBaryshe 2008).

Especially analogous to our model of TIR invasion and end module turnover are transposons of the Tn5053 family found in bacteria. These transposons are ‘res-site hunters’ that integrate into the resolution sites of other mobile elements, abolishing the functions of those resolution sites (Minakhina et al. 1999). Loss of resolution site function would be an evolutionary dead end for these mobile elements, but the Tn5053 family transposons encode their own resolution sites and recombinases, and most likely become an essential component of the elements they invade (Minakhina et al. 1999). The importance of replication and recombination for the maintenance of chromosomal ends likely makes these sites attractive targets for mobile element invasion. We suspect that element acquisition at chromosome ends and the domestication of such elements for end maintenance is likely a recurrent feature in evolution. In PLVs specifically, it may explain the high levels of genomic mosaicism.

## Conclusions

Here we describe the genomes of four novel PLVs, the first of such elements to be linked to poxviruses. The presence of three of these elements in EV occlusion bodies strongly suggests that they are hyperparasites of EVs. Symbiosis between these elements and EVs, along with their highly diverged capsid genes, suggests that our knowledge of PLV biology and diversity still remains fairly limited. Although several PLVs are linked with other viruses in the phylum *Nucleocytoviricota*, none have ever been found for the class *Chitovirales*, which includes the families *Poxviridae* and *Asfarviridae* (Koonin et al. 2020). Hence, our results suggest that hyperparasitic viruses may be a common feature of large DNA viruses in the *Nucleocytoviricota*. Given that many viruses in this phylum infect humans or agriculturally-important livestock, our results suggest that it may be possible to discover hyperparasitic viruses that could be useful for therapeutic or biocontrol applications.

Analysis of the four EV associated PLV genomes shows PLVs to be mosaic elements that have variable associations between different morphogenesis modules and the modules that facilitate both replication and integration. Recombination and integration modules appear linked to the PLV termini, suggesting that they might use the TIR as an origin of replication and recombination substrate for integration. We observe homology between these end modules and EV genes, including a number of helicases that show signs of mobility and appear to be linked to homing endonucleases. Given that homing endonucleases, recombination and homology to replication associated domains have been weaponized in virus-satellite conflicts (Barth et al. 2021; Nguyen et al. 2022; Barth et al. 2023), we expect the gene flow between EVs and PLVs to be a major driver of their evolution. We have also put forward a model that some PLV end modules are mobile, or at least have evolutionary roots as mobile elements.

## Materials and Methods

### Sequences analyzed

All genome sequences used in this work are publicly available. Major capsid protein sequences for previously recognized PLVs, apart from adintoviruses and MELD viruses, were obtained from Bellas et al. 2023 (Bellas et al. 2023). Major capsid sequences for adintoviruses and MELD viruses were obtained from Starrett et al. 2021 (Starrett et al. 2021) and Wallace et al. 2021. All other sequences analyzed appear in Table 1.

**Table 1.** Genomes used in this work and their associated NCBI accessions.

| **Name** | **Accession number** | **Notes** |
| --- | --- | --- |
| Mavirus (1) | KU052222 |  |
| Mavirus (2) | OX559518 |  |
| Mavirus (3) | OX560865 |  |
| Gezel-14T | NC_021333 |  |
| SFMaverick1 | BK011025 |  |
| SFMaverick2 | BK011026 |  |
| AHEV_PLV1 | KJ683044 |  |
| AHEV_PLV2 | KJ683045 |  |
| CBEV_PLV1 | KJ683046 |  |
| Ca_PLV1 | OX421890 | *Cetonia aurata*<br>chromosome 9<br>positions 23463350 to<br>23477421 |
| Adoxophyes honmai entomopoxvirus (AHEV) | HF679131 |  |
| Choristoneura biennis entomopoxvirus (CBEV) | NC_021248 |  |
| Choristoneura rosaceana entomopoxvirus (CREV) | NC_021249 |  |
| Mythimna separata entomopoxvirus (MySEV) | NC_021246 |  |
| Anomala cuprea entomopoxvirus (ACEV) | NC_023426 |  |
| Amsacta moorei entomopoxvirus (AMEV) | NC_002520 |  |
| Diachasmimorpha longicaudata entomopoxvirus (DIEPV) | KR095315 |  |
| Melanoplus sanguinipes entomopoxvirus (MSEV) | NC_001993 |  |
| Yalta virus isolate UA_Yal_14_16 (Yalta) | MT364305 |  |
| Heliothis virescens ascovirus 3g (HVAV_3g) | NC_044939 |  |
| Diadromus pulchellus ascovirus 4a (DPAV_4a) | NC_011335 |  |

### Gene function and structural prediction

Gene functional predictions were used using the HHPRED (Steinegger et al. 2019) tool hosted by the Max Planck Institute Bioinformatics toolkit (Zimmermann et al. 2018; Gabler et al. 2020).

Alphafold v2.2.2 (Jumper et al. 2021) was used to create monomeric models of all proteins in the genomes of AHEV_PLV1, AHEV_PLV2, and CBEV_PLV1. Twenty-five output structures were created for each model, ranked by confidence by Alphafold, the highest ranked of which was used for analysis. Proteins AJP61522, AJP61543, and AJP61551, respectively from AHEV_PLV1, AHEV PLV2, and CBEV_PLV1, each contained the folded beta sheets characteristic of a double jelly roll, and so were chosen as MCPs for each of the viruses. Top ranked models were assessed using SWISS-MODEL Structure Assessment and QMEAN(Benkert et al. 2011). AHEV_PLV1 had 80.28% Ramachandran favored residues with 5.54% outliers and a QMEAN Z-score of −4.38, AHEV_PLV2 had 85.44% favored residues with 4.67% outliers and a QMEAN Z-score of −4.84, and CBEV_PLV1 had 88.01% favored with 2.40% outliers and a QMEAN Z-score of −2.50.

### Phylogenetic tree construction

Protein sequence alignments were generated using Clustal Omega (Sievers et al. 2011; Sievers and Higgins 2018) using default parameters, and IQTree (Nguyen et al. 2015) was used to generate trees from the alignments, using ModelFinder Plus (Kalyaanamoorthy et al. 2017) for model selection and ultrafast bootstraps (Hoang et al. 2018). The best tree of 10 runs was selected. For the major capsid alignment, the alignment was trimmed using trimAl (Capella-Gutiérrez et al. 2009) prior to tree generation removing positions with a gap in 90% or more of sequences. For the D5-like helicase domains, full length proteins were manually trimmed prior to alignment based on locations of predicted domains and homologous regions. Trees were visualized using the iTOL webserver (Letunic and Bork 2021).

### Sequence similarity detection

Sequence similarity was detected through BLAST (BLAST+ 2.7.1) (Camacho et al. 2009) using the parameters word size 11, Match 4, Mismatch −5, gap costs existence 12 extension 8 for nucleotide sequences and word size 3 matrix BLOSUM62 and gap cost existence 11 extension 1 for peptide sequences. Alignments in Figure 4 B and C were generated using ClustalOmega (Sievers et al. 2011; Sievers and Higgins 2018) using default parameters.

## Acknowledgements

We thank Basil Arif for assistance collecting the original samples. We would also like to thank members of the Alward lab for useful discussion. We acknowledge use of the Virginia Tech Advanced Research Computing cluster for research performed in this study.

