## Supplementary figures and images for "Polinton-like Viruses Associated with Entomopoxviruses Provide Insight into Replicon Evolution"

### Figure S1

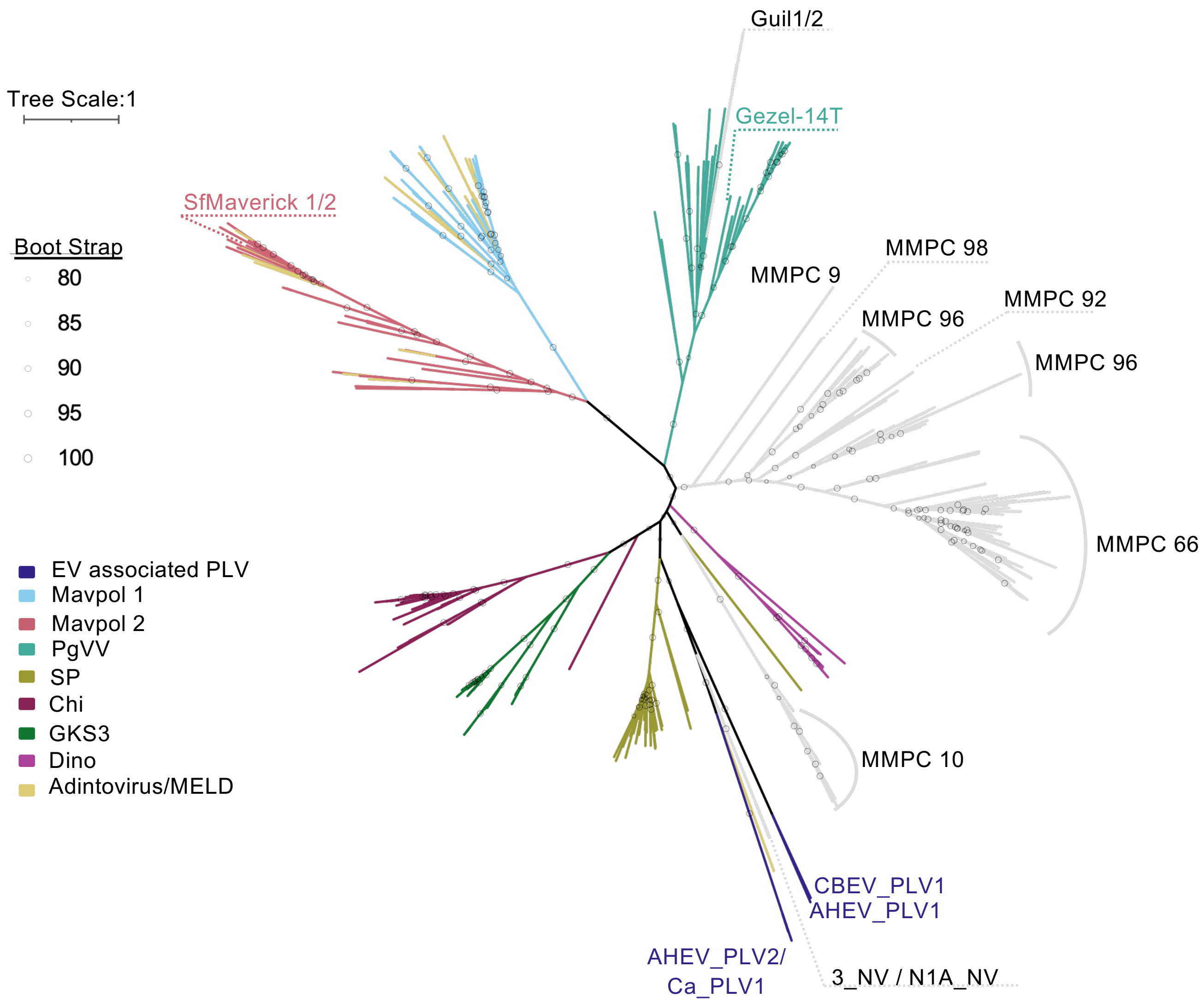
